# The tapejarid pterosaur *Tupandactylus imperator* from Crato Formation and the preservation of cranial integuments

**DOI:** 10.1101/2020.02.06.937458

**Authors:** Hebert Bruno Nascimento Campos, Edio-Ernst Kischlat

## Abstract

The group Tapejaridae forms a clade of toothless pterosaurs easily recognized by their premaxillary sagittal crests and particularly large nasoantorbital fenestrae. The tapejarids represent the most representative group of pterosaurs from the Lower Cretaceous Crato Formation of the Araripe Basin (Northeastern Brazil). The holotype of the large tapejarid *Tupandactylus imperator* Campos and Kellner, 1997 is known by two main slabs from the New Olinda Member of the Crato Formation, however, only one of the slabs containing the sagittally bipartite skull is referred to the holotype of *Tupandactylus imperator*, remain the counter-slab be properly described. The cotype is fragmented in several broken pieces and presents a significative number of cranial elements. A medial internasal septum completely preserved inside the nasoantorbital fenestra is reported for the first time for pterosaurs. The exceptional preservation of a collagenous septum and other integumentary structures visible in the cotype specimen is extremely rare and supports the concept of the unusual pattern of soft tissue observed in the fossils from the Crato Konservat-Lagerstätte, specially pterosaurs. Herein is presented the description of the cotype of *Tupandactylus imperator*, in complementation to the previously designated slab of the holotype of this tapejarid species. The occurrence of casques in pterosaurs is supported by comparative anatomy with the bird galliform *Pauxi* (Cracidae). Besides that, it is discussed on the skull with extravagant cranial crests of *Tupandactylus imperator* and the significance of the associated soft tissues and other cranial integuments, which indicates an expressive morphological and taxonomic diversity among the tapejarid pterosaurs.

## Introduction

The unusual preservation of the exceptionally well-preserved fossils from Lower Cretaceous Crato Konservat-Lagertätten of the Araripe Basin of Brazil (Varejão et al. 2019) already provided numerous exceptionally well-preserved pterosaur specimens, with a parcel associated to soft tissue, representing the most abundant amount of pterosaur skulls with exquisite preservation of soft cranial crest worldwide (Frey et al. 2003b; Pinheiro et al. 2019).

Tapejaridae is a clade of toothless pterosaurs recognized by their premaxillary crests and particularly large nasoantorbital fenestrae (Kellner 2004; Kellner and Campos 2014). The tapejarids represent the most representative group of pterosaurs from Crato Formation with three named species known so far: *Tupandactylus imperator* (Campos and Kellner 1997), known for some skulls (Pinheiro et al. 2011); “*Tapejara*” *navigans*, known by at least two skulls (Frey et al. 2003a) and an almost complete and undescribed skeleton; and *Aymberedactylus cearensis* based on a three-dimensional and almost complete mandible (Pêgas et al. 2016).

Based on the known specimens, the tapejarid *Tupandactylus imperator* is the largest species of the group. The holotype of *Tupandactylus imperator* is known by one skull sagittally bipartite embedded in two broken slabs typical of the Crato Formation, however, only one slab was originally described as the holotype (Campos and Kellner 1997), remain the counter-slab be properly studied. Herein is presented the description of the part of the skull of this bizarre tapejarid species contained in the counter-slab and is discussed about the associated soft tissue and preserved integuments.

## Geological Settings

The fossil-rich carbonate deposits of the Aptian Crato Formation of the Araripe Basin are one of the main Cretaceous Konservat-Lagerstätten of Gondwana (Varejão et al., 2019). The fossils from Crato Formation are confined to 8 meters thick, in the basal carbonate unit in the region of the city of Nova Olinda (Ceará, Northeastern Brazil), known as Nova Olinda Member (Martill, 1993).

The lithology of the Crato Formation is complex and differs from the other three units that compose the Araripe Group (Barbalha, Ipubi and Romualdo formations). It consists of laminated limestones and greenish shales sequences, which are also interbedded by sandstones and, occasionally, by thin layers of evaporitic minerals, predominantly gypsum (Martill, 1993). Mineralogically, the limestone of the Crato Formation is composed of fine-grained calcite crystals, with varying content of amorphous organic matter and siliciclastic minerals (Martill, 1993; Pinheiro et al., 2019).

The exceptionally well-preserved fossils from the Nova Olinda Member preserves organic structures such as integuments, that are observed only in a reduced number of pterosaur specimens (Frey et al., 2003; Unwin and Martill, 2007).

The tapejarid pterosaur *Tupandactylus imperator* comes from the lacustrine laminated limestones from Crato Formation. Although both slabs of the holotype of *Tupandactylus imperator* come from the Nova Olinda Member of the Crato Formation and belong the same individual, they have a distinct aspect of the lithology and colouring, which the slab has a yellowish colour while the counter-slab exhibits a beige colour.

## Description and Comparisons

### Skull

The skull of the holotype of the tapejarid *Tupandactylus imperator* (MCT 1622-R) is sagittally divided into two main slabs, which only one of the parts was shortly described (Campos and Kellner 1997). Almost cranial elements, as well as a portion of the soft tissue, also are preserved in the counter-slab, permitting that new morphological features and information on the cranial anatomy of this bizarre tapejarid be recognized.

The counter-slab containing the skull of the cotype of *Tupandactylus imperator* is displayed in six pieces, differing of the slab that is distributed on several fragmentary pieces. The calcareous slabs of the cotype specimen that contain the counter-part of the skull are more resistant, thicker and more compact than the slabs of the.

The skull of *Tupandactylus imperator* is proportionally low and anteroposteriorly elongated compared to other tapejarids. Except for the bone crests that extend posteriorly behind the skull, *Tapejara wellnhoferi* (Wellnhofer and Kellner 1991) and “*Tapejara*” *navigans* (Frey et al. 2003a) present a short skull compared to *Tupandactylus imperator*.

The skull roof is formed essentially by the combination of the frontals and parietals, as is typical of most pterosaurs. Compared to other tapejarids, *Tupandactylus imperator* presents the longer parieto-occipital crest. The tapejarid *Sinopterus benxiensis* from Jiufotang Formation of China also shows a long parieto-occipital crest (Lü et al. 2007), but it is directed upwards and not straight as in *Tupandactylus imperator*. The parietals in *Sinopterus benxiensis* extend posterodorsally by an angle of about 60° relative to the ventral margin of the skull (Lü et al., 2007).

The general morphology of the beak (pointed and toothless) was associated with a specific feeding strategy in tapejarids (Wellnhofer and Kellner, 1991). As is typical for *Tupandactylus imperator*, the rostral end is downturned, but the tip is missing. On the most rostral portion occurs a vestigial rhamphotheca and its internal aspect is internally visible. It is seen as displayed horizontally trabeculae and presents empty areas among them. On the most rostral portion is preserved a vestigial rhamphotheca (rhinotheca) located only on the rostral portion of the premaxilla. The internal aspect of the rhinotheca is visible. Internally rhinotheca is composed of trabecular bone displayed horizontally with present empty areas among them. The is seen as a thick whitish-coloured horny layer. The rhamphotheca represents an integumentary structure that is very rarely preserved in pterosaurs.

The nasoantorbital fenestra of the type specimen of *Tupandactylus imperator* is extremely large. It represents the largest cranial opening (250 mm length) and occupies approximately 60% of the overall length of the skull. Comparatively, the tapejarids “*Tapejara*” *navigans* and *Tapejara wellnhoferi* also present a large nasoantorbital fenestra that occupy more than 50% from the overall length of the skull (Frey et al., 2003a; Wellnhofer and Kellner, 1991). Compared to all other pterosaurs, the nasoantorbital fenestra of *Tupandactylus imperator* represents the larger structure known. The species of the toothed *Istiodactylus* also present large nasoantorbital fenestra, occupying more than half of the total skull length (Witton, 2012).

Based on the slab of the holotype (Campos and Kellner, 1997), the total length of the skull (from the tip of the premaxilla to the end of the parieto-occipital crest) is calculated in 800 mm and has an equal measurement for the high; the length from the tip of the premaxillary to the posterior part of the squamosal is equals to 420 mm; and the length of the most of the skull (the bone crests), is 670 mm.

Based on the dimensions of the skulls known, the proportions indicate that *Tupandactylus imperator* is the large tapejarid known (Table 1). Compared to “*Tapejara*” *navigans*, the skull length (from tip of the premaxillary to the posterior part of the squamosal) of *Tupandactylus imperator* is at least 10% large.

### Bone crests

Approximately half of the bone crests are preserved and are tabular in shape. Overall, no sutures can be identified. Following Pinheiro et al. (2011), in *Tupandactylus imperator*, the posterior extension of the premaxillae articulates as also are in direct contact with the nasals and frontoparietals, contrasting with the condition observed in *Tapejara wellnhoferi* and *Sinopterus dongi*, where there is a short space between these bones and the posterior extension of the premaxilla that runs parallel to the nasals and the frontoparietals. However, this condition also is noted in the cotype of *Tupandactylus imperator*.

The bone sagittal crests can be divided into the following different parts: The suprapremaxillary process, frontopremaxillary crest, fibrolamellar bone crest, premaxillary posterior extension, and parieto-occipital crest.

The anterior portion of the beak forms a ventro-dorsally tapered structure. It is filled with hollow structures. The fibrolamellar bone tissue is posteriorly joined with the frontopremaxillary crest and extends posteriorly above the premaxillary extension. Both structures, the fibrolamellar bone tissue, and the frontopremaxillary crest are best preserved in the slab of the type specimen that the noted in the cotype. The fibrolamellar bone tissue is distributed radially and forms an unusual part of the premaxillary crest.

The parieto-occipital crest extends beyond the final portion of the cranial box. Both, parieto-occipital crest and the premaxillary extension following parallel one each other.

### Soft-tissue

So far, the preserved nasal septum was reported exclusively on tapejarid pterosaurs from Crato Formation. The preservation of a medial internasal septum is extremely rare and was firstly reported for “*Tapejara*” *navigans* (Frey et al. 2003), which confidently mentioned the associate part of the soft structure inside the nasoantobital fenestra. But in the specimens known of “*Tapejara*” *navigans* only are preserved incomplete parts of the internal cranial septa.

The nasal septum is fully preserved on both slab and counter-slab of the holotype of *Tupandactylus imperator*. The septum occupies approximately 40% of the area of the nasoantorbital fenestra, and is seen as a region darker than the colour of the slab and is connected to the anterior portion of the nasoantorbital fenestra. It is possible for the presence of melanin in that structure.

In the cotype of *Tupandactylus imperator* the sagittal soft crest rests dorsally on top of the skull. It is bluish and differs markedly from the bony skull. However, only the proximal portion is preserved. No fibre is identifiable. Nevertheless, due to the appearance and the occurrence of granule concentration along the soft crest, it is probably the existence of melanin and melanosomes as noted in the crest of another skull of *Tupandactylus imperator* (Pinheiro et al., 2019).

The fibrous crest corresponds to 5/6 of the total lateral area of the skull in *Tupandactylus imperator* (Campos and Kellner, 1997), while in “*Tapejara*” *navigans* it occupies about 3/4 (Frey et al., 2003). In the cotype, the crest is lesser complete than in the type specimen.

The spine-like suprapremaxillary process is very thin and high. It sustain anteriorly the fibrous crest, and develops dorsally above a protuberant rostral structure derived of the premaxillary crest. The process inclines 15° posteriorly and represents the dorsal part of the rhinotheca. It has a whitish appearance as the rhinotheca, suggesting that is formed of the same keratinous component. As in “*Tapejara*” *navigans*, it is probably that the suprapremaxillary process represents merely an extension of the rhamphotheca.

### Taphonomy

In general, preservation of soft tissues in pterosaurs is relatively very rare in all the fossil deposits known. Body and cranial integumentary structures occur in several azhdarchoid pterosaurs from Crato Formation (Frey et al., 2003), and are formed by collagenous (e.g., soft cranial crest, wing membrane, internal cranial septa, pedal, and manual webbings) or keratin (e.g., keratinous sheaths, beak sheath, and integumentary fibres).

The exceptional preservation of integumentary structures with an original composition of collagen or keratin present in the pterosaur specimens from Crato Konservat-Lagerstätte is attributed to unusual mineralization, that permits that certain organic components degrade. It only occurs when the mineralization processes are faster than degradation (Briggs, 2003). It was suggested that benthic microorganisms may have promoted the protection of the fossils from Nova Olinda Member (Varejão et al., 2019).

According to Frey et al. (2003), the soft parts of the fibrous cranial crests, rhamphothecae and internasal septum present in the Crato pterosaurs are preserved as a goethite replacement, probably after pyrite in a process of pyritization, one of the main taphonomic processes involved in the exceptional preservation of soft-tissue.

The radial fibrolamellar bone tissue is located at the top of the premaxillary crest and develops posteriorly dorsal to the portion of the premaxillary extension back of the skull. According to Bantim (2017), the fibrous-parallel tissue is characterized by a rapidly depositing primary tissue. It represents a fast-developing support tissue, but also rigid enough to withstand conditions of constant mechanical stress. Based on the immaturity of the internal bone tissue, composed of a parallel fibrolamellar matrix of the premaxillary crest of the pterosaur *Maaradactylus kellneri* from Romualdo Formation, was indicated that it probably represents a sub-adult individual. Despite that, both pterosaurs present distinct external morphology of the premaxillary crest, similarly the same inference can be utilized for indicating the ontogenetic immaturity of *Tupandactylus imperator* (MCT 1622-R), representing also a sub-adult individual.

### Avian Comparative Anatomy

Like birds, the pterosaur beaks are formed by layered tissues, which contain a bony core and an outer keratin layer. The beak consists of bone (normally premaxillary and/or maxillary), vascular dermis and a modified germinal layer. The keratin layer of the beak is called the rhamphotheca, with the rhinotheca covering the upper beak and the gnathotheca covering the lower beak (O’Malley, 2005). In birds and pterosaurs, the keratin layer forms hard-cornified tissues and consists of living cells that keratinize towards the outer side of the beak, leaving a hard, cornified, dead keratin layer at the outer surface of the beak.

The casques usually constitute a well-defined and prominent cranial protuberance Mayr (2018). According to Naish and Perron (2016), casques primarily represent novel hypertrophy of frontal bones that are involved in the casque formation. Casques and cranial crests are widespread in pterosaurs in very different shapes. The tapejarid *Tupandactylus imperator* exhibit an extravagant complex of bone crests on the top of the skull, which represents a unique configuration of the skull. At least three bones are involved in the formation of the cranial crests of *Tupandactylus imperator*: premaxillae, distributed rostrally; and occiput and parietals that are distributed back of the skull.

Morphologically the cranial crests of *Tupandactylus imperator* are quite different from any bird. However, the beak can be analogically compared to bird casques. According to Mayr (2018), cranial bony protuberances of birds often exhibit a continuous ontogenetic growth and in several species of these groups, their development is sex-dependent. In the species of the galliform *Pauxi* (Cracidae) both sexes develop a bulb-like protuberance at the base of the culmen called dorsal ridge (Mayr, 2018). In most of the protuberance of the upper beak of *Pauxi* occurs an only slightly cornified, bluish integument (Vaurie, 1968).

The avian skull exhibits a large structural diversity, in that bony cranial protuberances can be present. In birds, the protuberances of the upper beak usually develop from the nasal and premaxillary bones. As in *Tupandactylus imperator*, the cranial crest of *Pauxi* quit developing rostrally growing up. In *Pauxi*, the crest can occupy the same height of the skull but differs from *Tupandactylus imperator* in that the premaxillary crests are divided into four distinct types: the massive crest, the laminar crest and the posterior prolongation that sustain a median fibrous crest. The bulbs of the species of *Pauxi* exhibit slightly cornified skin (Mayr, 2018). The same integument may have covered part of the cranial crests of *Tupandactylus imperator*.

Due to the proportions, the nasoantorbital fenestra noted in the tapejarid *Tupandactylus imperator* can be regarded as not analogous to any species of related groups, once that this developed cranial structure is comparatively larger than any bird, or non-avian theropod known. Hypothetically, apart from the morphological implications (e.g., occupying a significative area of the skull), it is probable that the function of the large nasoantorbital fenestra in pterosaurs, as tapejarids and istiodactylids, is involved in a powerful olfactory capacity, permitting the catching long-distance prey odours.

The bird *Cathartes aura* (Falconiformes, Cathartidae) presents at least two cranial features that are analogy with the pterosaur *Tupandactylus imperator*: The downturned edentulous beak and a proportionally large narial opening. According to Grigg et al. (2017), a sensitive sense of smell allowed the turkey vulture to colonize biomes that are suboptimal for scavenging birds and become the most widespread vulture species in the world. A hypothetical inference would enable pterosaurs with large nasoantorbital fenestra a significative eco-physiological advantage and adaptive distribution in relation to other predatory/scavenging pterosaurs.

## Discussion

Apart the high ingroup diversity, the tapejarid pterosaurs form a clade with high proportional divergence with a range of known species and with a large distribution worldwide, which includes *Sinopterus benxiensis* Lü et al. (2007), *Sinopterus corollatus* Lü et al. (2016), *Sinopterus dongi* Wang and Zhou (2003), *Sinopterus lingyuanensis* Lü et al (2016) and *Sinopterus atavismus* Lü et al. (2016) from Lower Cretaceous of China (Jiufotang Formation, Aptian); *Caiuajara dobruskii* Manzig et al. (2014) from Upper Cretaceous of Brazil (Goio-Erê Formation, Turonian); *Tupandactylus imperator* Campos and Kellne (1997), *Aymberedactylus cearensis* Pêgas et al. (2016) and “*Tapejara*” *navigans* Frey et al. (2003a) from Lower Cretaceous of Brazil (Crato Formation, Aptian); *Tapejara wellnhoferi* Kellner (1989) from Lower Cretaceous of Brazil (Romualdo Formation, Albian); and *Europejara olcadesorum* Vullo et al. (2012) from Lower Cretaceous of Spain (Calizas de La Huergina Formation, Barremian).

Although there are pterosaurs three-dimensionally preserved from Crato Formation, they represent an exception (e.g., Pêgas et al. 2016). Frequently, fossils from Crato Formation are compacted and are embedded in a single calcareous slab, or are divided on two distinct limestones divided sagittally, with one aspect of the specimen complementing the another, as observed in a basal serpentiform snake (e.g., Martill et al., 2015).

The tapejarid *Tupandactylus imperator* was the first-named pterosaur for the Crato Formation (Campos and Kellner 1997), and it is the species with most numerous numbers of specimens known, with at least five skulls housed in public and private collection (Pinheiro et al. 2011). Furthermore, it is the most representative pterosaur in relation to the soft tissue association, with all the known specimens associated with exceptionally well-preserved soft crest (Frey et al. 2003b; Pinheiro et al. 2019). This fact indicates that the *Konservat-Lagertätte* Crato Formation is the most important fossil deposit for that type of unusual soft-tissue preservation.

The cotype of *Tupandactylus imperator* described here represents an additional specimen from the same brood as the original type specimen defined as the main slab of the skull preserved in the doubly distributed calcareous concretions. This is the case of the specimens referred to as the holotype of *Tupandactylus imperator*, which represents both parts of a single specimen. The cotype is properly available on a public collection (Museu de Ciências da Terra, Rio de Janeiro, Brazil), and despite the skull is less complete that the part of the type species, it present diagnostic characters for *Tupandactylus imperator* (an extremely large soft-tissue median cranial crest supported anteriorly by a spine-like suprapremaxillary process caudally inclined and an anteriorly projecting convex blade on the premaxillae) and tapejarid features (e.g., downturned beak, very large nasoantorbital fenestra, pear-shaped orbit and premaxillary crest extending behind the skull). Which permit confidentially assigns the skull preserved on the counter-slab to *Tupandactylus imperator*.

The large nasoantorbital fenestra represents the large cranial opening and is an important tapejarid feature. It occupies about 60% of the overall length of the cranial skeleton of *Tupandactylus imperator*. Together with the species of *Istiodactylus* (Witton, 2012), *Tupandactylus imperator* exhibits the large nasoantorbital fenestra among all pterosaurs known. The physiological function of the proportionally large nasoantorbital fenestra in pterosaurs is unknown. Anatomically, the nasoantorbital fenestra to decrease the total weight of the skull, permitting a higher performance in the full head balance.

## Conclusions

The counter-slab of the holotype of the tapejarid pterosaur *Tupandactylus imperator* from the Early Cretaceous Crato Formation is described. The specimen represents a cotype in complementation to the designated type specimen. It preserves part of the cranial rhamphotheca (rhinotheca) and other integumentary structures of the skull. The cotype adds new morphological information on *Tupandactylus imperator*, supporting the knowledge on the extravagant cranial morphology and complexity of the configuration of the crests of this taxon. The cranial crests of tapejarids show a significative diversity in shapes and proportions. They are originated from soft and osseous tissue and the high variation observed indicates an important taxonomic diversity in this pterosaur group during the Cretaceous. Finally, based on the exceptionally well-preserved specimens known, the *Konservat Lagertätte* Crato Formation of Brazil represents the most important fossil deposit of this type of unusual preservation of soft tissue of cranial structures in pterosaurs as crests, internasal septa, and keratinous rhamphothecae.

**Figure 1.**
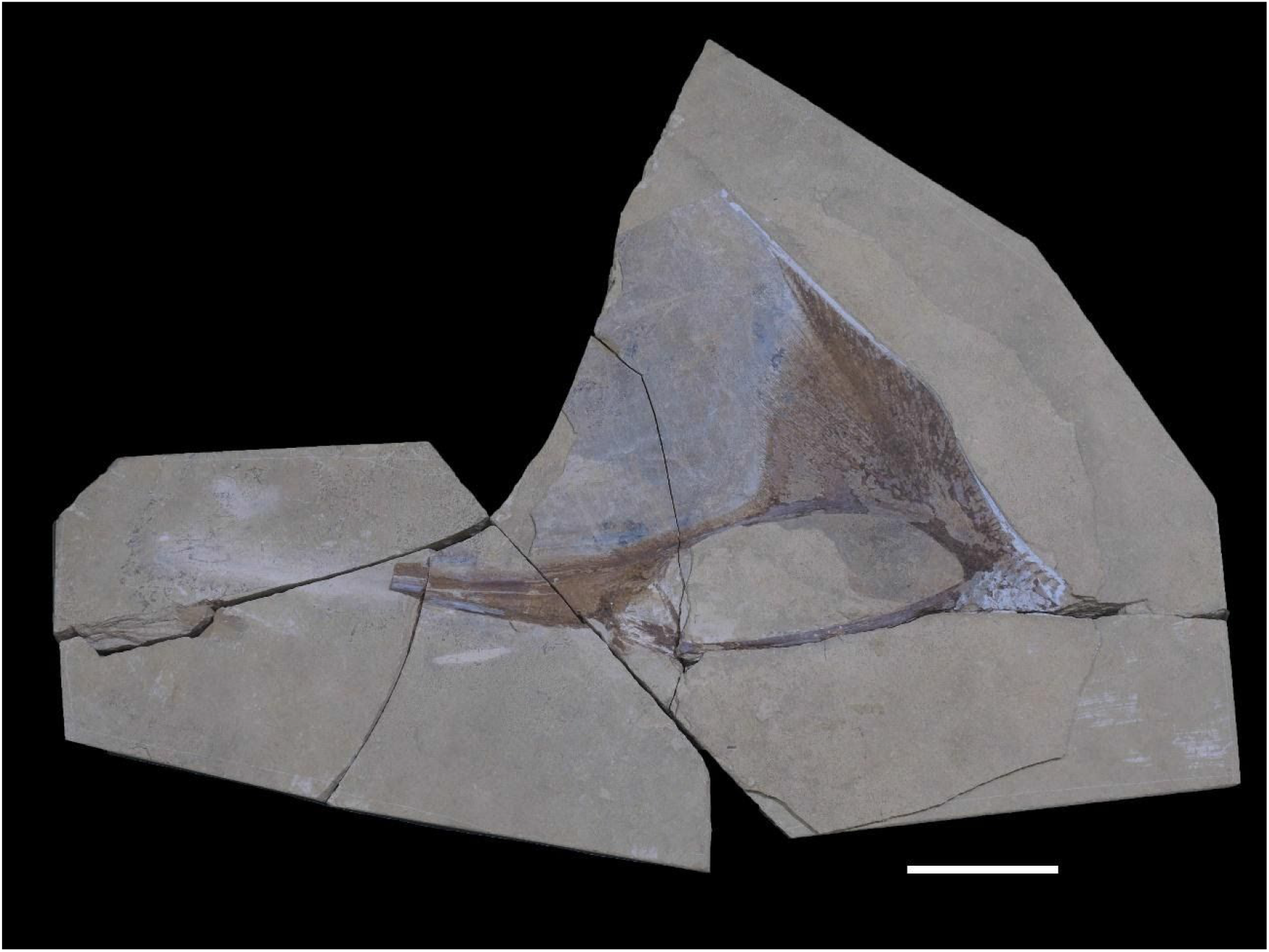
Cotype of *Tupandactylus imperator* (MCT 1622-R). Scale bar equals to 100 mm.

**Figure 2.**
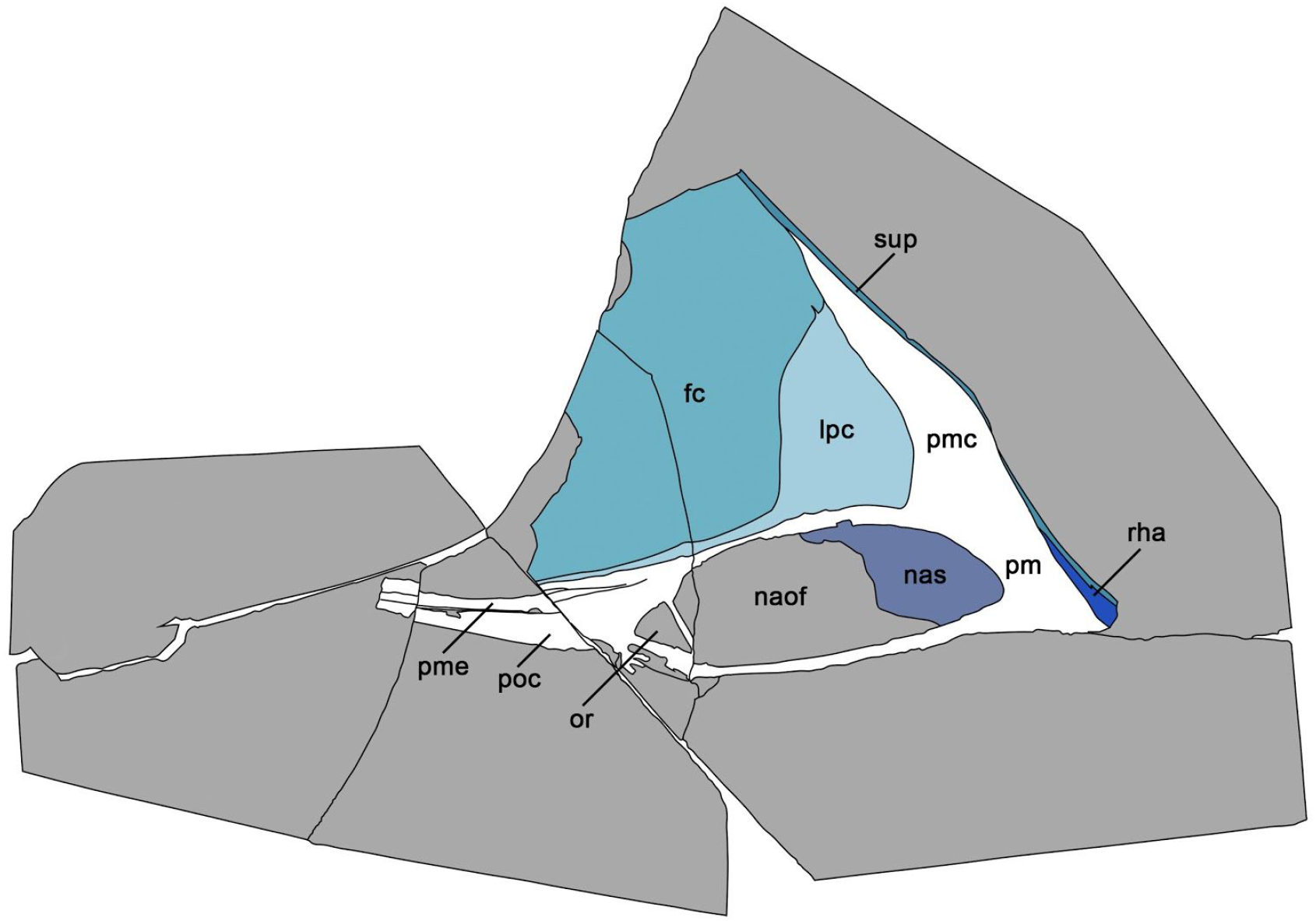
Cotype of *Tupandactylus imperator* (MCT 1622-R), with marked separated elements of the cranial skeleton and integuments. Abbreviations: fc, fibrous crest; lamellar premaxillary crest; naof, nasoantorbital fenestra; nas, internasal septum; or, orbit; pmc, premaxillary crest; pm, premaxillary; pme, premaxillary extension; poc, parieto-occipital crest; rha, rhamphotheca; sup, suprapremaxillary crest.

**Figure 3.**
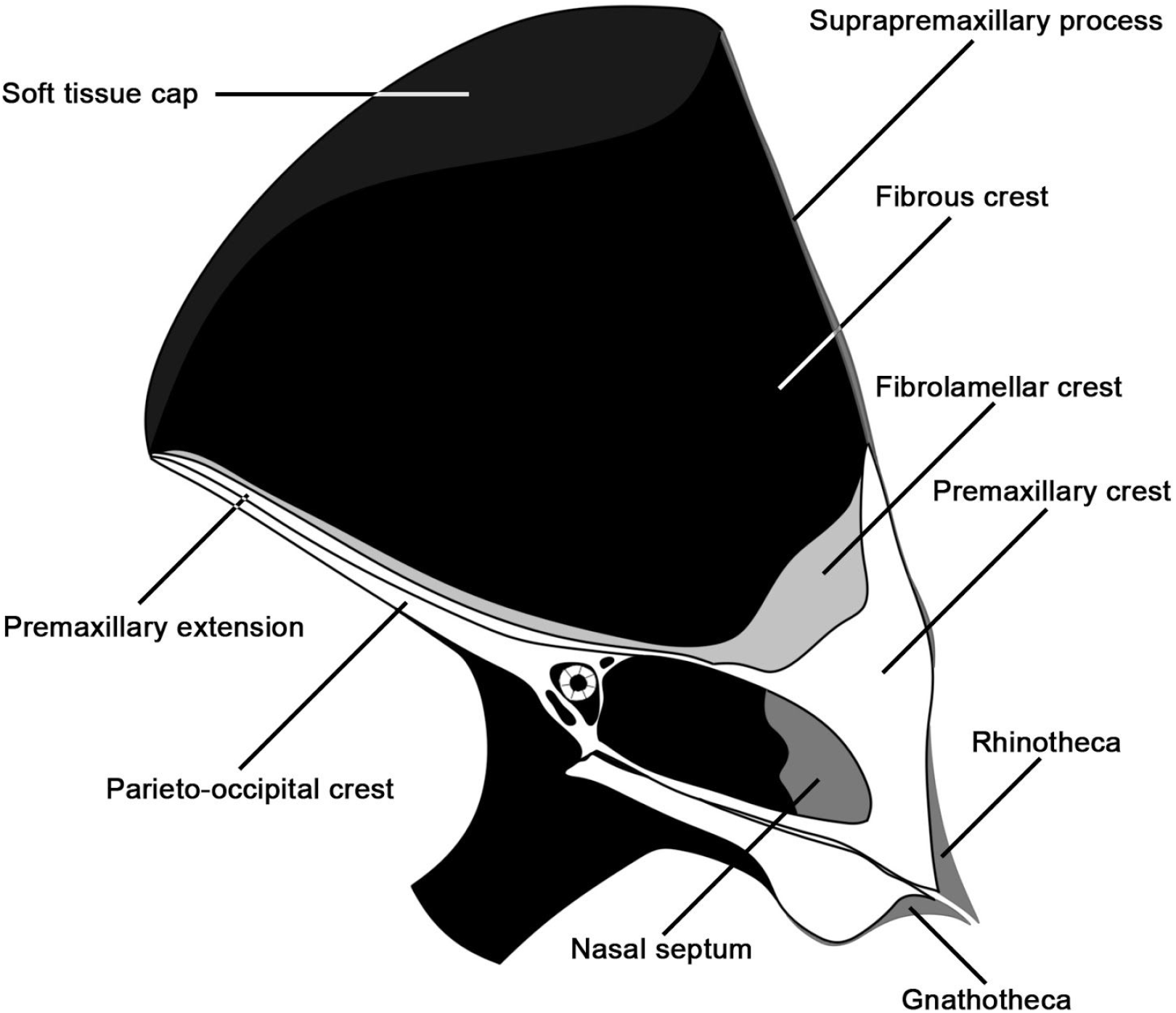
Schematic reconstruction of *Tupandactylus imperator* indicating the main cranial structures. Based on the hypothesis of Frey et al. (2003) and specimens of Campos and Kellner (1997), Pinheiro et al. (2011) and this paper.

**Figure 4.**
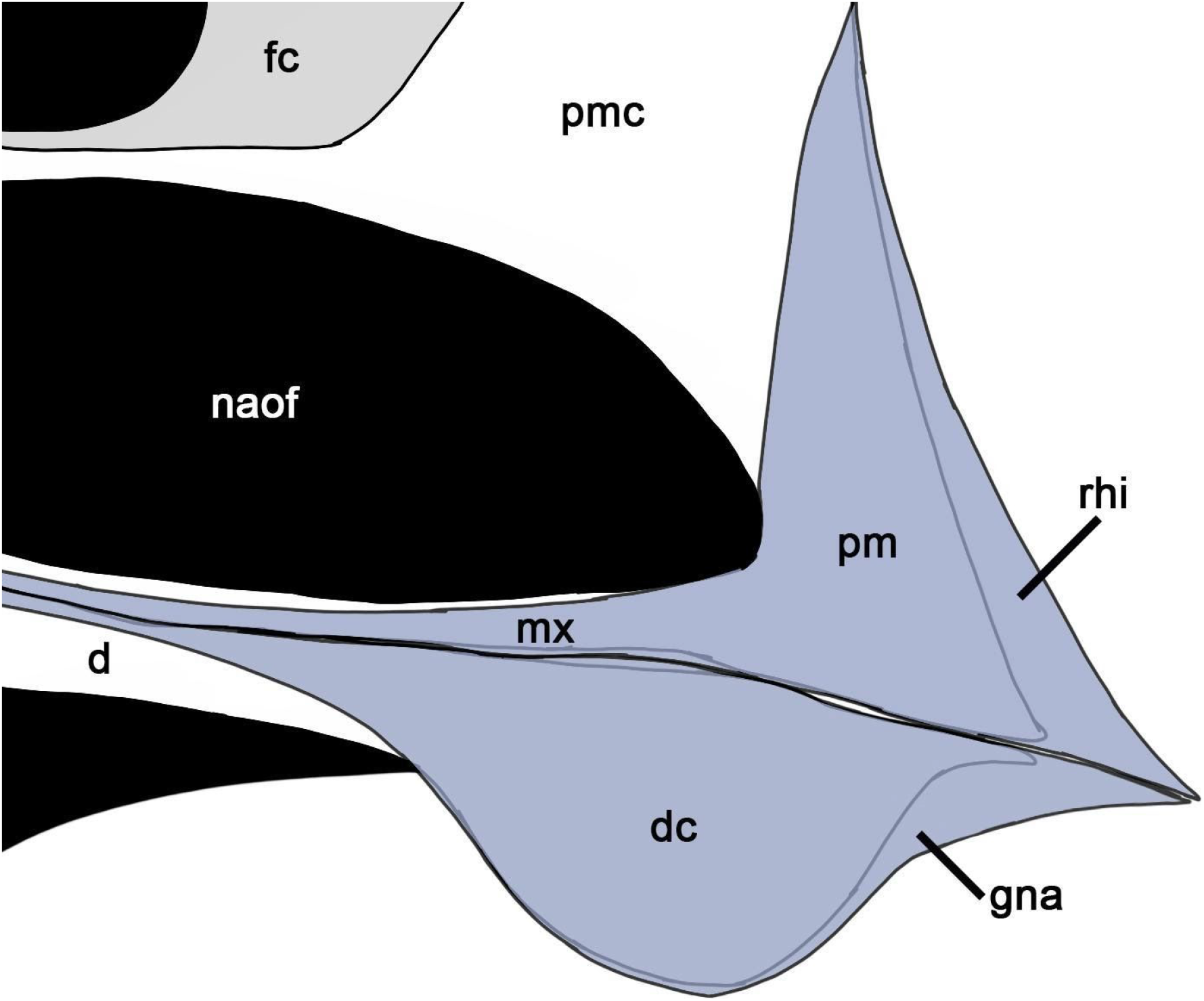
Schematic representation of the hypothetical basal portion of the multi-layered beak of *Tupandactylus imperator* covered by a rhamphotheca. Based on the specimens known (Campos and Kellner, 1997; Pinheiro et al., 2011, and this paper). Abbreviations: d, dentary; dc, dentary crest; fc, fibrous crest; gna, gnathotheca, mx, maxillary, pm, premaxillary, pmc, premaxillary crest; rhinotheca.

## Acknowledgements

Thanks to Prof. Dr. Eberhard “Dino” Frey (Naturkunde Museum Karlsruhe, Karlsruhe, Germany) for permission to access the specimens of tapejarids under your care. Thanks to Dr. Rafael Costa da Silva (Museum of Earth Sciences, Geological Survey of Brazil, Rio de Janeiro, Brazil) for permission to publish the specimen described here. Thanks to Prof. Dr. Alexander W. A. Kellner for permission to access the pterosaur specimens deposited at the National Museum (UFRJ, Rio de Janeiro, Brazil). Thanks to Dr. André Veldmeijer (The American University in Cairo, Cairo, Egypt) to share their photographs of Araripe pterosaurs. Finally, thanks to Nicholas Gardner (Potomac State College, Keyser, West Virginia, USA) for the comments on the early version of this manuscript and Dean Schnabel (Germany) for the use of their illustrations. Thanks to the Geological Survey of Brazil (CPRM) for support to this study. This study was partially funded by the Brazilian National Council for Scientific and Technological Development (CNPq) [Process Number: 290025/2015-0].

